# Applications of T-cell receptor specificity annotation models for quality control and immunomonitoring in adoptive T-cell therapies

**DOI:** 10.64898/2026.01.26.701708

**Authors:** Max Van Houcke, Sander Wuyts, Tom Bosschaerts, Johanna Chiffelle, Aymeric Auger, George Coukos, Alexandre Harari, Pieter Meysman

## Abstract

Adoptive cell therapy (ACT) with tumor-infiltrating lymphocytes (TIL) is a powerful candidate immunotherapy. However, the personalized production process that underlies its potential also introduces an element of stochasticity, where the actual target of the TILs is variable across patients and protocols, and remains largely unknown. In this letter, we describe the application of T-cell receptor (TCR) sequencing in combination with computational tools for annotating epitope-specificity of the TCR repertoire of TIL products as a potent quality control. We highlight the potential of this approach by demonstrating the tumor antigen-specificity of TCR clusters within the TIL product *in silico*, and validating these responses *in vitro*. We also demonstrate screening for off-cancer reactivity against viral antigens, and immune monitoring of identified TCR clones.

**Highlights:**

- T-cell receptor sequencing combined with epitope annotation tools are viable for quality control of TIL products by identifying tumor-specific clones.
- We showed the additional utility of screening for off-cancer reactivity, such as viral antigens.
- The identified tumor-specific TCR clones can be tracked in vivo providing a means of immune monitoring.

## 1. Introduction

T-cells form the basis of the next revolutionary generation of cancer therapeutics. Their central role in immune responses drives groundbreaking innovations from T-cell based therapeutics to novel diagnostics, which underscores the importance of studying T-cells and T-cell responses. Adoptive Cell Therapies (ACT) allow the use of an individual’s own T-cells to combat cancer. Tumor-infiltrating lymphocytes (TILs) are T-cells resident within tumors, representing a heterogeneous collection of T-cells that recognize a broad range of cancer-associated antigens. TILs derived from resected tumors can be cultured and expanded in the absence of the immunosuppressive environment of the tumor. TIL therapy typically consists of surgically removing a tumor or tumor biopsy, isolating naturally occurring TILs from the tumor, and expanding and reinvigorating them to a large number (> 1000-fold expansion to generate billions of TILs) using an expansion protocol [1, 2]. The final TIL product is reinfused into the patient after a preparatory chemotherapy (lymphodepletion) to boost their ability to attach tumor. Treatment with first-generation TIL products has resulted in deep and durable responses in melanoma patients [3]. As a key example, the clinical results of a single-arm study were sufficient for accelerated conditional FDA approval of a TIL product for the treatment of refractory metastatic melanoma [4]. Clinical application of adoptive T-cell therapy with TILs is now practiced at many sites around the world, and an approximate 40-50% objective response rate is consistently observed across different sites for patients with melanoma [5].

The next frontier of TIL adoptive cell therapy is focused on improving response rates in melanoma and expanding the applicability of TILs to other solid tumor types. A key success of TIL therapy relies on the quality, specificity, and number of pre-existing TILs [6]. Next-generation TIL products need to be optimized for phenotype and tumor specificity. Novel manufacturing protocols that drive the rapid expansion of tumor-reactive CD4+ and CD8+ T-cells within TIL cultures, actively selecting tumor-reactive T-cells or genetically modifying TILs to express optimal phenotypes, are currently being explored [3,7]. A final TIL infusion product is assumed to have a high frequency and high variability of tumor-reactive T-cells, a crucial parameter for clinical success. Indeed, a study in refractory metastatic melanoma patients revealed that treatment with TIL products with demonstrable T-cells reactive against tumor epitopes was correlated with higher response rates [8]. However, it is well documented that TILs can also be reactive to common viral antigens [9]. As some TILs are not tumour-reactive, so-called bystander T-cells, identifying which of the many T-cells in the tumour microenvironment and the final TIL product recognise tumour antigens is a long-standing quest. This is especially problematic as virus-specific T-cells have been shown to have a higher target affinity than tumor-specific T-cells [10], which can cause an undesirable skewing during the manufacturing process. Once a TIL product has been manufactured, it is therefore not guaranteed to only contain tumor-reactive T-cells and requires a second quality control step. Potency assessment of TIL products currently lags the clinical efficacy of these therapies, hindering further clinical trials and subsequent commercialization of TIL therapies [11].

Multi-omic sequencing approaches are being used to understand the optimal TIL phenotype associated with response and design more potent and persistent TIL products. T-cell receptor (TCR) sequencing has emerged as an accurate and reproducible method for profiling TIL products [12,13]. TCR sequencing is already performed routinely at different timepoints during the manufacturing process by clinical sites and companies developing ACTs, in order to monitor and evaluate clonal expansion and product diversity. In addition, TCR sequencing on peripheral blood mononuclear cells (PBMCs) or tumor samples post ACT enables tracking the persistence of relevant TIL clones *in vivo*. The TCR plays a crucial role in T-cell functionality as it defines the specificity of the immune response by recognizing peptide sequences (epitopes) derived from pathogens, cancer cells or autoantigens. Epitopes are presented in the context of human leukocyte antigen (HLA) molecules on the surface of antigen-presenting cells, and recognised by TCRs for effective immune responses. As the TCR defines the epitope-specificity of a T-cell, it can be used to evaluate the antigen-specificity of the TIL product. However, the high TCR diversity and HLA polymorphisms make the identification of tumor-reactive T-cells from their TCR sequence a non-trivial problem and practically unfeasible with *in vitro* techniques. Computational methods have emerged to solve the annotation problem, and recent benchmarks have shown that the current state-of-the-art has achieved the necessary performance for practical applications [14].

In this paper, we evaluate the potential of TCR-epitope annotation to characterize TIL ACT products. We demonstrate that the epitope-specificity of T-cell clusters in the TIL product can be determined *in silico*, both for tumor-associated and viral antigens. We show validation of the annotations made by the algorithms through *in vitro* experiments with a transgenic T-cell line and epitope-specific stimulation experiments. In addition, we demonstrate that identified tumor-specific T-cells can be tracked *in vivo* using bulk TCR sequencing on PBMCs, as a means of immune monitoring after ACT.

## 2. Methodology

### 2.1 Bulk TCR sequencing data of tumor, TIL products and PBMC repertoires

All TCR sequencing data are identical to those described in Chiffelle *et al*., 2024 [6] studying tumor-reactive T cell clonotype dynamics underlying clinical response to TIL therapy in melanoma, accessible under GEO accession number GSE229861. The results in this study correspond with those from the subject labelled ‘patient 11’ within the original study.

In brief, we deeply interrogated T cell populations and TCR repertoires from patients who received TIL therapy in a phase-I clinical trial (NCT03475134). TILs were expanded *in vitro* from patients’ tumor deposits resected by surgery under high doses of interleukin-2 (IL-2) in the GMP manufacturing facilities of the Centre Hospitalier Universitaire Vaudois (CHUV) under specific conditions previously described [15]. Some TILs of the infusion product, as well as resected tumor samples, were cryopreserved for translational studies including bulk TCR sequencing. A blood sample was obtained at the same time as tumor surgery and at a minimum of 30 days after adoptive TIL transfer, and peripheral blood mononuclear cells (PBMCs) were cryopreserved. Bulk TCR sequencing analyses were performed using SEQTR, an in-house developed sensitive and quantitative assay for TCR repertoire analyses, as described previously [16].

### 2.2 Computational analyses

TCR clonotype files designated as “TCRbeta” were retrieved from GEO accession number GSE229861. These clonotype files were preprocessed by the original authors using ad hoc Perl scripts to: (i) pool all TCR sequences coding for the same protein sequence; (ii) filter out all out-frame sequences; (iii) determine the abundance of each distinct TCR sequence.

TCRs with a single read were not considered for the analysis [6]. All clonotype files were directly used as input for subsequent analysis, without any other preprocessing steps. Pielou’s Evenness of these repertoires is defined as the Shannon entropy divided by the maximum score, namely the natural logarithm of the number of unique TCRs.

To evaluate TCR similarity within TIL products, bulk TIL TCR repertoires were clustered with ClusTCR, a python interface for rapid clustering of large sets of CDR3 sequences with unknown antigen specificity (version 1.0.0) [17] and visualized with python-igraph (version 1.0.0) (Figure 1). ClusTCR groups TCRs together based on their CDR3 beta amino acid sequences and a Hamming distance of one (a maximum of one amino acid difference between any two connected TCRs).

**Figure 1.**
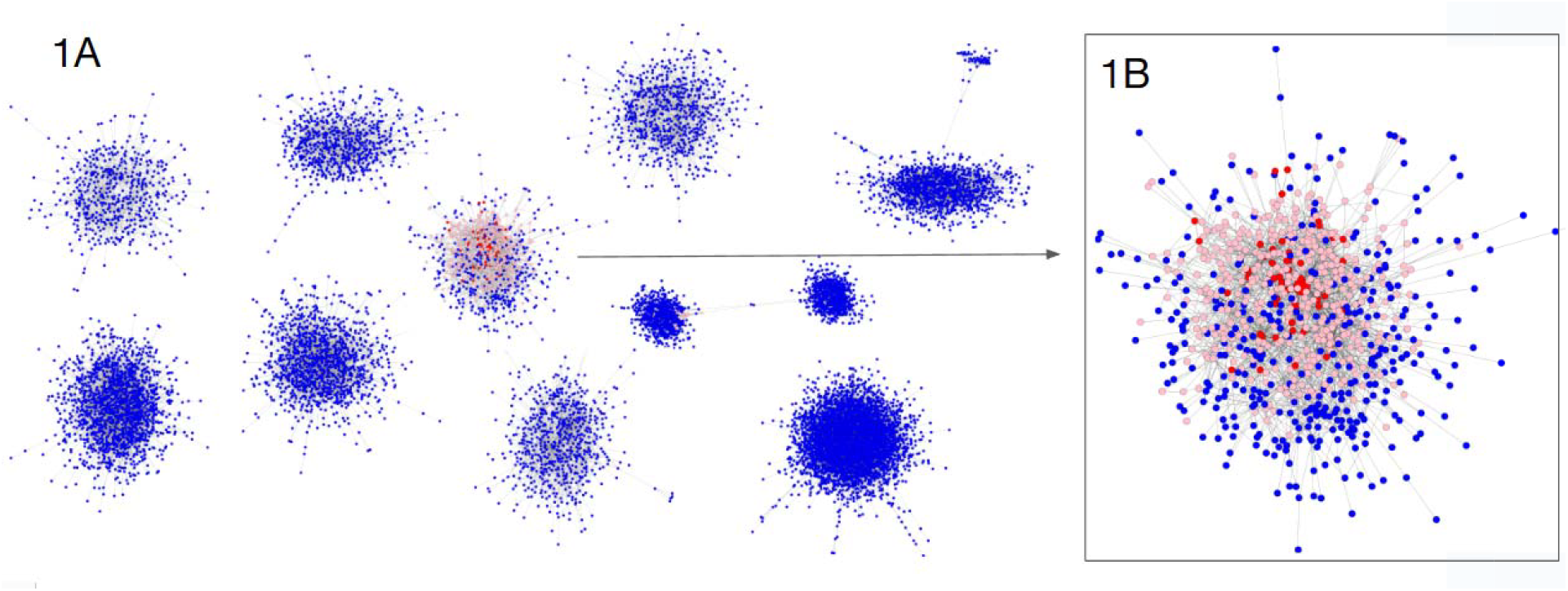
A. T-cell receptor network of the TIL product of patient 11, where each node constitutes a unique beta TCR, and each edge denotes high sequence similarity between two beta TCRs. B. Annotations with ImmuneWatch DETECT, T-cells annotated to be tumor-reactive against the MelanA ELAGIGILTV epitope are depicted in red (highly confident predictions above cut-off score 0.2) or pink (medium confidence predictions with DETECT score between 0.15 and 0.2).

Identification of epitope-specificity of TCRs was performed using ImmuneWatch DETECT, a command-line solution for epitope-TCR annotations (Version 1.4.3) [18]. ImmuneWatch DETECT is a seen epitope model consisting of machine learning models and a training database of epitope-TCR pairs (IMWdb), and has been shown to reliably annotate TCRs for epitopes covered in its training database [14]. IMWdb contains training data for over 2000 epitopes, including those for viral and melanoma antigens. The recommended scoring thresholds of 0.2 (high confidence) and 0.15 - 0.2 (medium confidence) were used for analysis of epitope specificity [18].

### 2.3 TCR cloning and tumor reactivity validation

Methods to interrogate antitumor reactivity are described in Chiffelle *et al*. [6] and Schmidt *et al*. [19]. Both bulk TCR sequencing and single cell sequencing datasets are available under GEO accession number GSE229861. Beta T-cell receptors of interest identified in the bulk TCR sequencing were screened for in the single cell sequencing datasets to find their matching alpha TCR chain. After matching, TCR alpha-beta (TCRab) pairs were cloned for functional validation. These TCRab pairs were expressed in reporter Jurkat cell line (TCR/CD3 Jurkat-luc cells (NFAT), Promega, stably transduced with human CD8alpha/beta and TCRalpha/beta CRISPR-KO). The T-cells were activated and expanded before being transfected with the TCRab chain RNA using an electroporation system. Following transfection, the modified cells were co-cultured with autologous PBMCs and stimulated with specific peptides (control irrelevant peptide, MelanA peptide ELAGIGILTV, no peptide), or treated with TransAct as a positive control (a polymer conjugated to recombinant CD3 and CD28 agonists for activation of T cells). For the reporter Jurkat cells, NFAT pathway activation upon antigen recognition was quantified via the Bio-Glo Luciferase assay (Promega).

## 3. Results

### 3.1 Characterisation of a TIL product with TCR clustering and epitope-TCR annotations

Bulk TCR sequencing data from a TIL infusion product sample, as described in Chiffelle *et al*. [6], were analysed for TCR similarity and for antigen-specificity. Clustering of bulk beta TCR sequencing data of the TIL product uncovered 40,127 TCRs with a Pielou’s Evenness score of 0.395, indicating a highly oligoclonal population. A selection of the 10 most abundant TCR clusters in this TIL product is depicted in Figure 1, representing about 70% of the total TCR repertoire in the TIL product. Annotation of epitope-specificity with ImmuneWatch DETECT revealed that there was a single cluster with a dominant signal. The majority of T-cells in this cluster were annotated to be reactive to the immunodominant ELAGIGILTV epitope of MelanA, a well-known melanoma-associated antigen (Figure 1).

### 3.2 Evaluating the presence of virus-specific TCRs in TCR repertoires

Human immune repertoires contain many virus-specific TCRs, and TILs have also been reported to be virus-reactive. Most of the epitope-TCR training data for epitope annotation models, such as ImmuneWatch DETECT, is for viral epitopes (approximately 50% of 2200 training data epitopes in the IMWdb database). To assess the presence of virus-specific TCRs in the TIL product, we analysed available TCR samples with ImmuneWatch DETECT and filtered on TCRs scoring positive for viral epitopes. The final ACT product showed a low percentage of virus-specific TCRs (0.02% of TCRs scoring positive for any virus with DETECT), in particular compared to the hits in other samples such as PBMCs (ranging between 0.36% and 1.52%) and baseline tumor sample (0.70%), indicating little to no viral antigen-specific expansion in the TIL product. The MelanA-specific annotations took up 9.8% of the TIL repertoire, overall showing a relatively higher content of tumor-specific TCRs and lower content of virus-specific TCRs compared to the baseline samples (Figure 2), denoting a successful tumor antigen-specific lymphocyte expansion.

**Figure 2.**
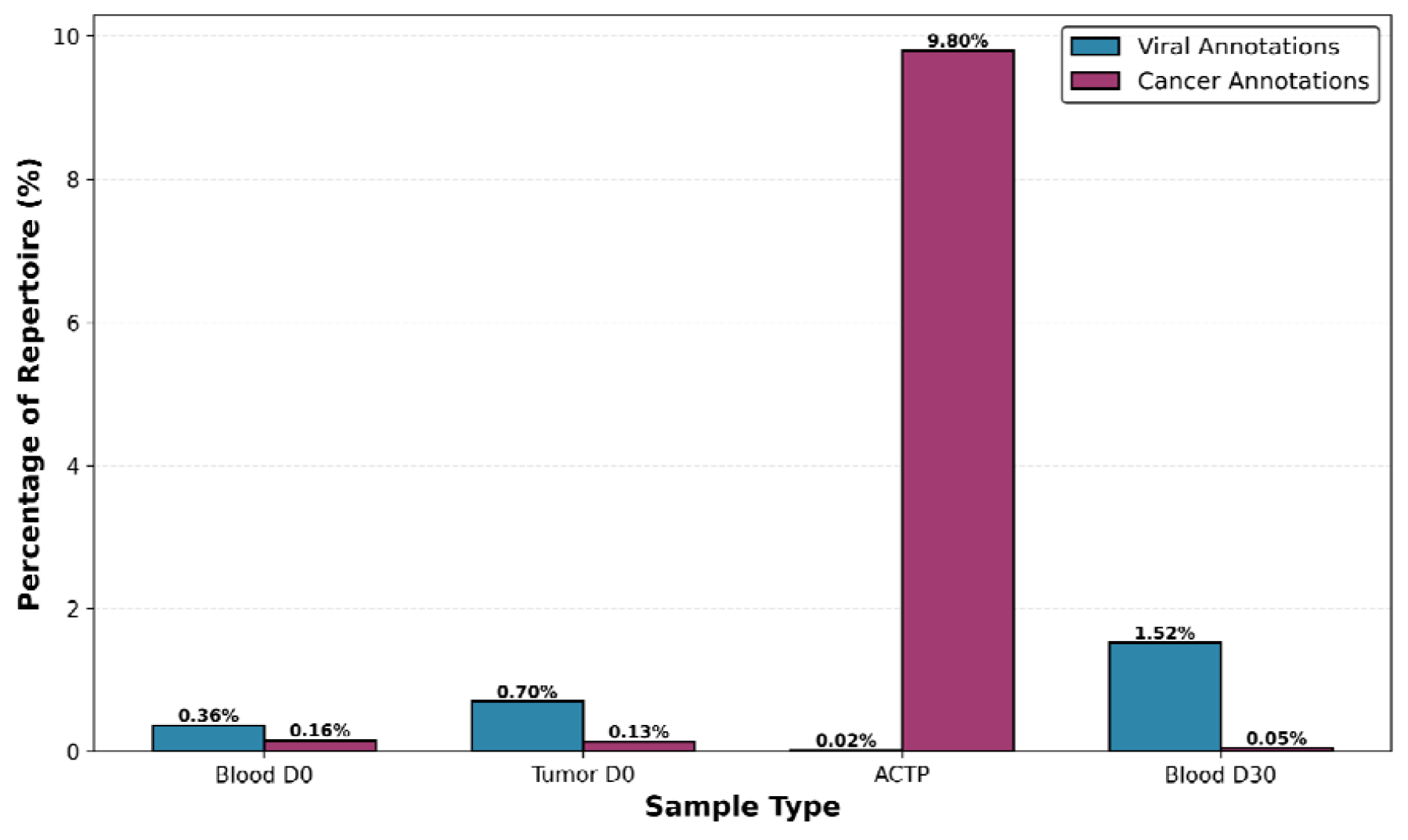
Viral and MelanA-specific TCR annotation levels. Virus-specific TCRs are almost non-existent in the ACT product (0.02%) while present in PBMC and tumor TCR repertoires. The ACT product is enriched in MelanA-specific TCRs.

### 3.3 Validation of the epitope-specificity of annotated MelanA-specific TCRs

To confirm the antigen-specificity of predicted TCRs from the cluster highlighted in Figure 1B, we selected three distinct TCRs with high frequencies and reliable tumor-antigen annotation in the TIL product for *in vitro* functional validation of their putative epitope target ELAGIGILTV. The most dominant TCRb chain consisted of almost 10% of the total repertoire (Table 1). Based on single-cell TCR sequencing data, every TCRb chain of interest could be mapped and paired with an alpha chain. The paired sequences from Table 1 were cloned in a reporter cell line, co-cultured with autologous PBMCs, and stimulated with the ELAGIGILTV peptide, alongside positive and negative controls. There was a clear and specific T-cell activation upon stimulation with the ELAGIGILTV peptide for all selected TCRs (Figure 3). These experiments demonstrate that TCR clusters identified by ImmuneWatch DETECT were indeed reactive against the ELAGIGILTV epitope.

**Table 1:**
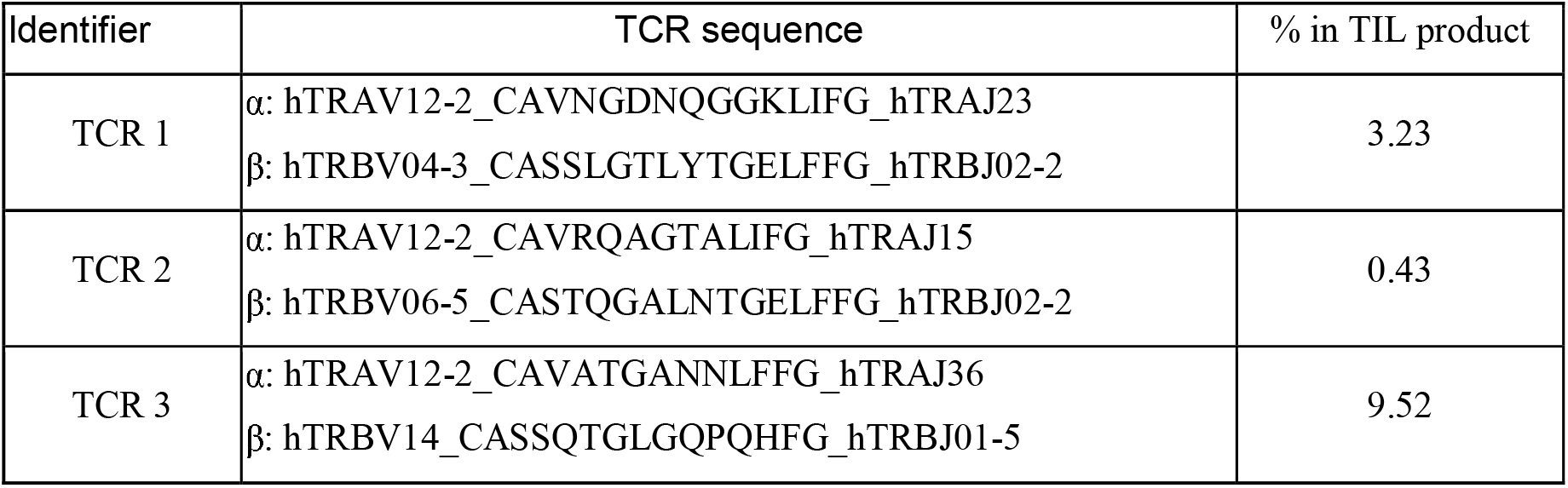
Paired alpha and beta TCR sequences selected for validation, based on ImmuneWatch DETECT prediction of MelanA-specificity. Percentage of each of these clones within the entire TIL product is also shown.

| Identifier | TCR sequence | % in TIL product |
| --- | --- | --- |
| TCR 1 | $\alpha$: hTRAV12-2_CAVNGDNQGGKLIFG_hTRAJ23<br>$\beta$: hTRBV04-3_CASSLGTLYTGELFFG_hTRBJ02-2 | 3.23 |
| TCR 2 | $\alpha$: hTRAV12-2_CAVRQAGTALIFG_hTRAJ15<br>$\beta$: hTRBV06-5_CASTQGALNTGELFFG_hTRBJ02-2 | 0.43 |
| TCR 3 | $\alpha$: hTRAV12-2_CAVATGANNLFFG_hTRAJ36<br>$\beta$: hTRBV14_CASSQTGLGQPQHFG_hTRBJ01-5 | 9.52 |

**Figure 3.**
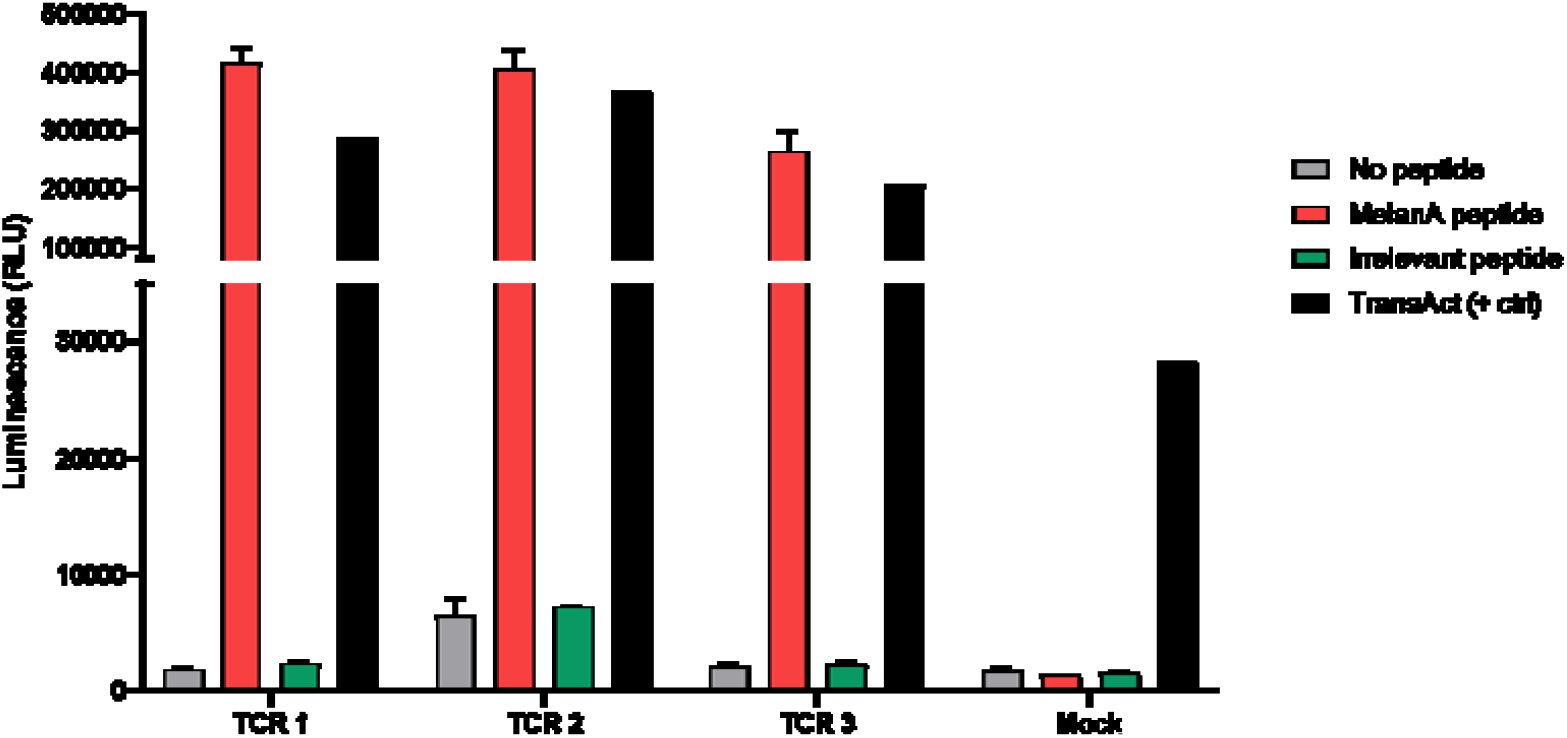
Overview of TCR functional validation experiment for TCR1, TCR2, and TCR3 from patient 11 TIL product. In the absence of any peptide as a negative control (grey), the ELAGIGILTV peptide (red), a control peptide (green) and TransAct as a positive control (black). Irrelevant mock TCRs were also transfected and used as negative control.

### 3.4 Immunomonitoring of MelanA-specific TCRs before and after ACT

Alongside data from the TIL infusion product, we analysed bulk TCR repertoires from pre- and post-treatment blood from the same patient. Our objectives were to i) identify individual TCRs from the MelanA-specific TIL cluster in peripheral blood samples, and ii) track these individual TCR clones before and after ACT. The pre-ACT peripheral repertoire contained 713,832 unique TCRs and the post-ACT sample 386,115 TCRs. The clonotypes identified within the cluster of ELAGIGILTV-specific T-cells could be tracked by their TCR sequence in these samples to characterise their dynamics following treatment (Figure 4). Several MelanA-specific TCR clones could be identified both pre- and post-ACT in this patient. In this instance, the MelanA-specific cluster decreased in frequency in this patient based on the peripheral TCR repertoires (Figure 4). It is noteworthy that the patient had a non-responder status after ACT.

**Figure 4.**
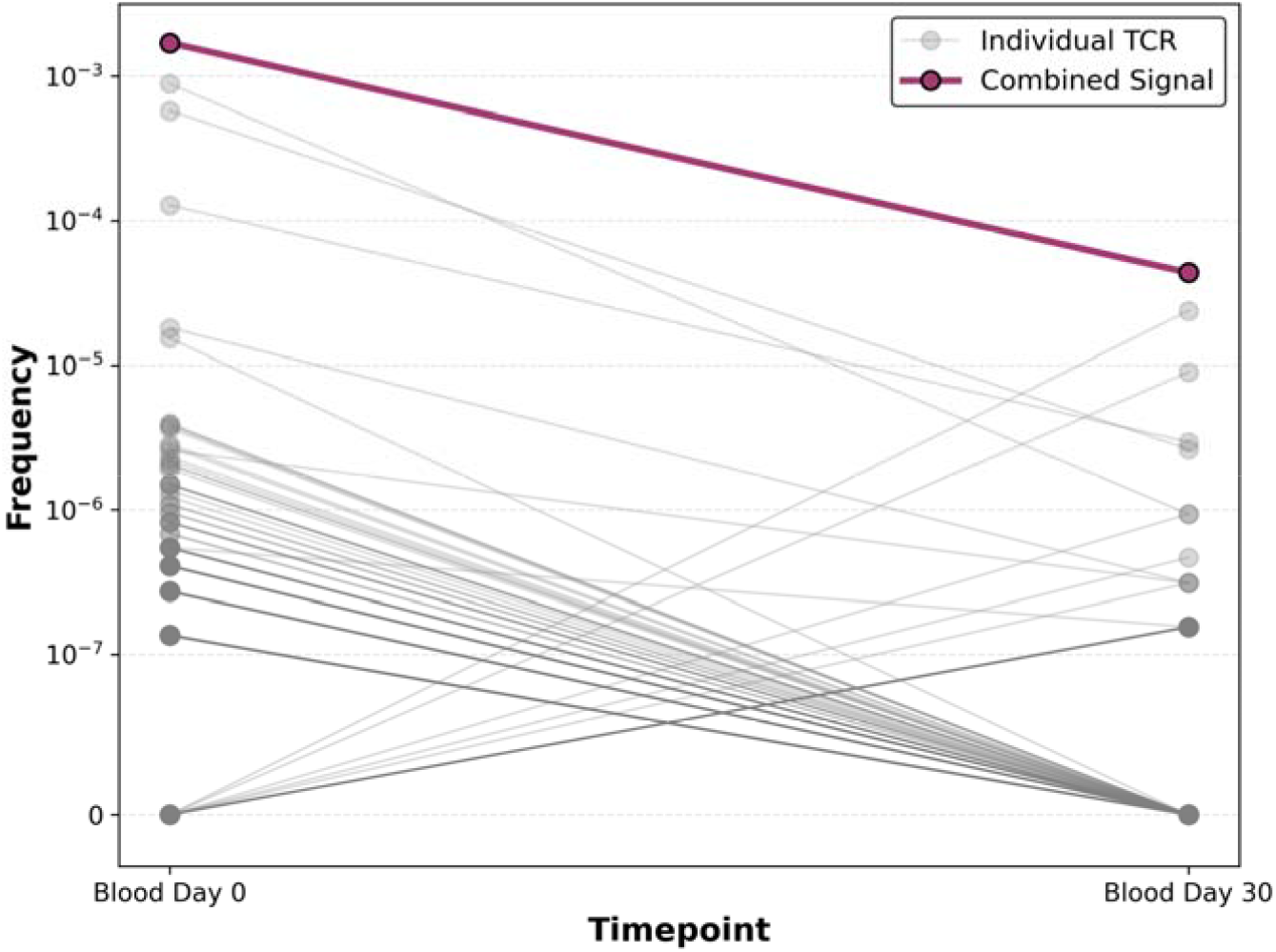
Repertoire frequency dynamics of MelanA-specific TCRs in PBMC samples before and after ACT. Dynamics of each TCR clonotype has been plotted as a distinct grey line. In bold purple, the combined signal of all MelanA-specific TCRs.

## 4. Discussion

TCR sequencing is a rapidly growing application with significant potential for advancing our understanding of the immune system and more effectively developing new therapies for various diseases. However, the inability to clinically interpret TCR sequencing data currently limits its value. The astronomical potential sequence diversity of epitopes (20M+ human 10mers alone [20]) and TCRs (10^61^ in the total theoretical space [21] and around 10^10^ in a single human [22]) has presented a barrier to computational prediction of TCR epitope specificity. However, knowledge of the target epitopes of TCRs will have a huge impact on the development and application of novel immunotherapies, vaccines, and diagnostics. This is a major bottleneck in immunotherapy development, particularly in the field of TIL-ACT, as the antigen-specificity of TIL products is currently mostly unknown.

Current T-cell receptor annotation algorithms are capable of identifying epitope-specific T-cells in full repertoires with high accuracy, however they remain restricted to a limited set of epitopes for which TCR training data is available. All available technologies for TCR-epitope annotation, including tcrdist3 [23], TCRex [24], NetTCR [25], MixTCRpred [26] and ImmuneWatch DETECT are “seen epitope” algorithms, for which an epitope of interest already has examples of TCRs binding it in the database [27]. As confirmed by recent IMMREP benchmarks organised by top-tier international academic consortia [14], seen epitope models perform well when training data is available.

Current performance of epitope-TCR annotation solutions can already add significant value to the evaluation of TIL-ACT products. A straightforward example is the detection of virus-specific TCRs, as a plethora of TCR data for viral epitopes is already available in the public domain and actively used as training data. Detecting viral T-cell clones at different timepoints of product development could be integrated into current TCR data analysis pipelines for TIL product characterisation and quality control. We also show that with available tumor-specific epitope-TCR training data, such as for common tumor-associated antigens like MelanA, identifying tumor-specific TILs *in silico* is feasible. Moreover, by cloning a selected set of TCRs and demonstrating their specificity for the MelanA epitope *in vitro*, we have validated the predictions made by the epitope-TCR annotation algorithms.

Immune monitoring of patients following ACT or other immunotherapies is an important aspect of clinical evaluation and biomarker development, often bottlenecks in translational sciences. In-depth analysis of TCR repertoires at single cell and bulk level, both for ACT product and pre/post ACT clinical samples, offers opportunities to simultaneously evaluate therapy-, virus-, or disease-specific responses. We demonstrate here the identification and tracking of MelanA-specific TCRs both in the TIL product and across PBMC samples.

Future efforts should focus on generating more and better-quality training data. Predicting TCR specificity is not an algorithmic challenge, but a data challenge [28]. Existing TCR-epitope training data is still too sparse and biased towards common HLAs and immunodominant viral epitopes. To further characterize TIL TCR repertoires in the absence of epitope-specific data, such as for neo-epitopes, additional analyses could be considered. Combining clustering of TCR with epitope motif generation and HLA predictions can significantly reduce the search space for epitope matches for a TCR cluster of interest. Screening of public TCR repertoires databases (such as iReceptor [29], or TIL-specific repertoires) could also be valuable to determine if TCR clones in TIL products have already been reported in similar clinical conditions.

## 5. Conclusion

In this study, we have demonstrated the potential use of a combination of sensitive bulk TCR sequencing and T-cell epitope annotation solutions such as ImmuneWatch DETECT as a viable assessment tool for TIL ACT therapies, both for TIL product characterisation and monitoring relevant T-cell clones *in vivo*. While the capacity is currently limited by the restricted set of immunogenic epitopes that can be annotated, concerted efforts across academia and industry to improve the underlying annotation capabilities will result in a potent quality assessment tool and potential companion diagnostic for ACT in the near future.

## Acknowledgements

Dr. Raphaël Genolet who performed bulk TCR sequencing and Alexandra Michel and Christophe Sauvage who screened TCRs for their antigenic specificity.

